# Direct Modulation Index: a measure of phase amplitude coupling for neurophysiology data

**DOI:** 10.1101/2022.02.07.479380

**Authors:** Maximilian Scherer, Tianlu Wang, Robert Guggenberger, Luka Milosevic, Alireza Gharabaghi

## Abstract

Neural communication across different spatial and temporal scales is a topic of great interest in clinical and basic science. Phase-amplitude coupling (PAC) has attracted particular interest due to its functional role in a wide range of cognitive and motor functions. Here, we introduce a novel measure termed the direct modulation index (dMI). Based on the classical modulation index, dMI provides an estimate of PAC that is bound to an absolute interval between 0 and +1, resistant against noise, and reliable even for small amounts of data. To highlight the properties of this newly-proposed measure, we evaluated dMI by comparing it to the classical modulation index, mean vector length, and phase-locking value using simulated data. We ascertained that dMI provides a more accurate estimate of PAC than the existing methods and that is resilient to varying noise levels and signal lengths. As such, dMI permits a reliable investigation of PAC, which may reveal insights crucial to our understanding of functional brain architecture in key contexts such as behaviour and cognition. A Python toolbox that implements dMI and other measures of PAC is freely available at https://github.com/neurophysiological-analysis/FiNN.

**Highlights:**

- Neural coupling measures are sensitive to higher harmonics of the target oscillation.
- dMI achieves frequency-specificity by sine-fitting the phase-amplitude histogram.
- Increased robustness to noise and signal duration in comparison to other measures.
- dMI allows for reliable estimation of phase-amplitude coupling.

## Introduction

For a number of years, the investigation of the communication between structures across different spatial and temporal scales has been a major area of interest in the field of cognitive and motor neuroscience (Siems et al., 2016; Siems and Siegel, 2020). In particular, a growing body of research regards phase-amplitude coupling (PAC) as a phenomenon reflective of a multi-frequency communication mode across and within neural structures (Canolty and Knight, 2010; Jensen and Colgin, 2007). The level of PAC between neural structures is quantified by the degree to which the phase of a low-frequency neural oscillation reflects the shape of the amplitude of a high-frequency oscillation (Bragin et al., 1995; Lakatos et al., 2005). Several studies have reported a close relationship between PAC of high-gamma amplitude with alpha phase and behavioural performance on cognitive, motor, and sensory tasks (e.g., Schroeder and Lakatos, 2009; Voytek et al., 2010; Yanagisawa et al., 2012).

Given the large interest in PAC, many methods have been developed to provide an accurate quantification of this cross-frequency neural communication (Tort et al., 2010). The methods established include the modulation index (MI; Tort et al., 2008), mean vector length (MVL; Canolty et al., 2006) and phase locking value (PLV; Mormann et al., 2005). Albeit these methods were successful in revealing relevant brain-behaviour relationships (e.g., Canolty and Knight, 2010; Penny et al., 2008), they share two main limitations. First, none of these methods provide a bounded output measure, which prohibits the interpretation of absolute PAC values across different studies (Hülsemann et al., 2019; Tort et al., 2010). Second, some methods may erroneously detect high PAC values at harmonic multiples of the frequency of the enveloped amplitude signal (e.g., Giehl et al., 2021; Kramer et al., 2008), as suggested by the findings of the present investigation.

This work introduces the direct modulation index (dMI) as a novel measure of PAC which aims to circumvent the limitations of the aforementioned methods. The dMI is a bound variation of the MI as introduced by Tort and colleagues (2008). It facilitates the interpretation of absolute PAC values, thus not only increasing the comparability of PAC values between different studies, but also allowing for interpretations beyond relative changes in PAC. Furthermore, dMI is highly sensitive to the target frequency only, and therefore avoids the pitfall of assuming significant PAC changes at harmonic frequencies to be actual findings. In the next section, we begin with a description of the proposed measure, followed by an illustration of its performance in comparison to a selection of established connectivity methods on simulated data.

## Methods

### Direct Modulation Index (dMI)

With the modulation index as its first step, the dMI shares the calculation of a phase-amplitude histogram (Tort et al., 2008). Briefly, using the Hilbert transform, a phase-amplitude histogram is constructed by extracting the phase of the low-frequency signal and the amplitude of the high-frequency signal. A composite time series is then constructed from the phase of the low-frequency signal and the amplitude of the high-frequency signal, and the mean amplitude is calculated across phase bins. From here, the calculation of dMI deviates from that of the classical MI. First, the phase-amplitude histogram is normalized to produce a more robust fit in the following step. To ensure that it is more resistant towards outliers, the 25^th^ and 75^th^ percentiles – as opposed to the minimum and maximum values – are chosen as reference points for the normalization. This results in a normalized histogram which is loosely bound to the interval [0, 1]. Second, instead of scoring the PAC values from entropy, a sinusoid is fitted to the normalized histogram. We selected a sinusoidal function because the phase-amplitude histogram of two signals with an ideal PAC relationship was observed to default to a sinusoid shape. Fitting a sinusoid through the phase-amplitude histogram renders the measure highly sensitive to the targeted frequency only, whereas the entropy metric is also sensitive to the harmonic frequencies. Finally, an error value is calculated by taking the squared difference between the height of each individual phase bin and the amplitude of the sinusoidal fit at the corresponding bin. The errors are averaged across phase bins, capped at 1, and then subtracted from 1 to arrive at the dMI. This final step simplifies the interpretation of the PAC estimate, as values approaching zero indicate low PAC, while values approaching 1 indicate high PAC

### Validation data

dMI was evaluated in comparison to the following PAC methods: MI (Tort et al., 2008; 2010), MVL (Canolty et al., 2006), and PLV (Mormann et al., 2005). The reader may refer to the corresponding articles for a description of the evaluated measures. dMI and the aforementioned PAC methods were evaluated using simulated data. As opposed to using experimental data – where it is unclear whether any detected PAC at harmonic frequencies is reflective of a true coupling – simulated data enabled us to absolutely determine any signal properties, including the confirmable absence of PAC at the higher harmonic frequencies. We generated a high frequency signal of 200 Hz with an amplitude that was modulated by a 10 Hz oscillation (Figure 1A), and several low frequency signals of 2, 5, 10, 15, 20, 25, and 30 Hz (Figure 1B). Each signal had a duration of 30 seconds and a sampling rate of 1000 Hz. A high PAC value was anticipated for the 10 Hz low frequency signal, but not for the other low frequency signals (Figure 1C).

**Figure 1.**
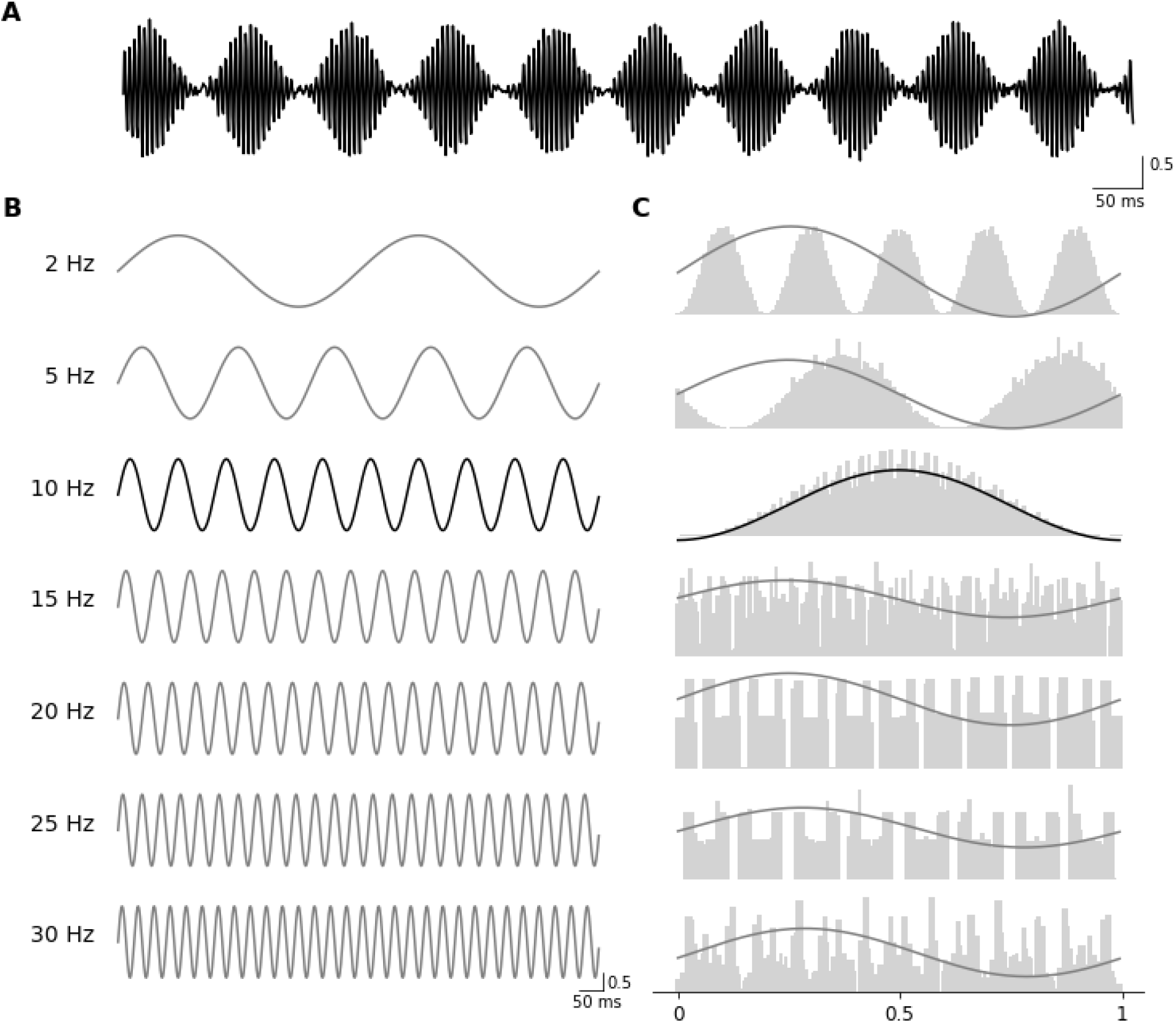
Simulated signals and calculation of direct modulation index (dMI). (**A**) High-frequency signal with an amplitude (in arbitrary units) that was modulated by a low-frequency signal. (**B**) Low-frequency signals used in the validation of dMI. The 10 Hz signal in the third row represents the low-frequency signal that modulates the high-frequency signal in (A). (**C**) Histograms of the amplitude of the high-frequency signal from (A) binned according to the phase of the low-frequency signals in (B) are shown in grey. The y-axis shows the mean amplitude in arbitrary units, the x-axis shows the phase bins in normalized units. The best sine fit is plotted over the histogram. The goodness of fit is calculated from the difference between the fitted sine and the height of the individual histogram bins. In the event of a good fit, as with the 10 Hz low-frequency signal in the third row, the sine fit lines up with the phase-amplitude histogram.

To test the performance of each PAC method in the presence of noise, Gaussian noise, with amplitudes that are 0%, 25%, 50%, 100%, or 150% of the amplitude of the signal, was introduced. To additionally investigate the combined effects of noise level and signal duration on each PAC method, the analysis was repeated with a signal length of 300 s. Finally, the performance of each method with varying signal durations was tested by capping the signals at durations of 500, 600, 700, 800, 900, and 1000 ms. In this evaluation, the amplitude of the Gaussian noise was fixed at 33% of the signal amplitude.

The scripts for this analysis can be found at https://github.com/VoodooCode14/dmi.

## Results

Figure 2A shows the PAC estimates obtained using dMI, MI, MVL, and PLV. A consistent, single peak at 10 Hz can be observed for dMI, with a PAC value of 0.94. PAC estimates from MVL showed elevated peaks at 5, 10, and 25 Hz, with values of 0.001, 0.24, and 0.006, respectively. PLV-based estimates showed consistent peaks at 10 Hz with a PAC value of 0.54. However, at frequencies of 5, 20, 25, and 30 Hz, elevated PAC values of 0.006, 0.13, 0.006, and 0.06, respectively also occurred. Finally, MI-based estimates showed elevated PAC values of 3.84 at 10 Hz, as well as values of 3.08 and 3.29 at the lower frequencies of 2 Hz and 5 Hz, respectively.

**Figure 2.**
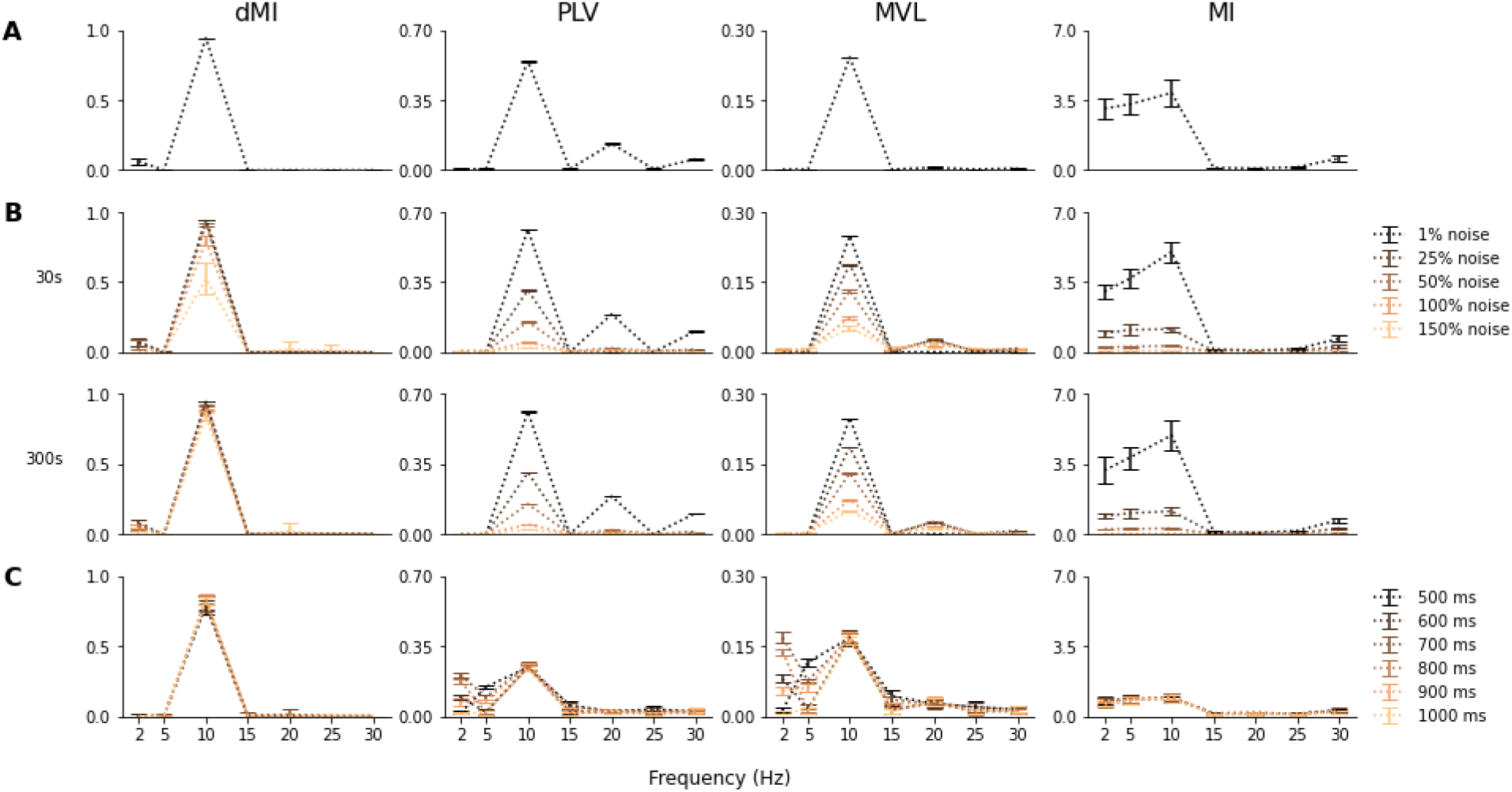
Comparison of PAC measures in simulated data. The columns show the performance of direct modulation index (dMI), phase-locking value (PLV), mean vector length (MVL), and the classical modulation index (MI). The y-axes show the PAC estimates in arbitrary units (mean and standard deviation). (**A**) PAC estimates on simulated signals of 30 s and 5% Gaussian noise. (**B**) PAC estimates on simulated signals of 30 s (top row) and 300 s (bottom row) with varying levels of Gaussian noise. (**C**) PAC estimates on simulated signals of varying durations and 33% Gaussian noise.

Figure 2B shows the effect of varying levels of Gaussian noise on the PAC estimates of the investigated measures for signal lengths of 30 s and 300 s. A general reduction in PAC values can be observed for increasing noise levels. PAC estimates based on MI and PLV decreased most (61% and 51%, respectively) in the presence of weak noise (25% of the signal amplitude). By comparison, the MVL-based estimates decreased by 25%, and the dMI-based estimates decreased by 0.29% only. Also, at Gaussian noise levels, dMI drop rates were consistently lower than those of the other evaluated measures. Furthermore, increasing the signal length to 300 s increased the robustness against noise for the dMI-based PAC estimates, but not for the other measures (Figure 1B, bottom row). Finally, the erroneously elevated PAC values of MI and PLV were also present in the analysis across different levels of Gaussian noise.

Figure 2C shows the effect of signal length on the PAC estimates of the measures investigated. The results were similar to those reported in the previous investigation of the effect of noise. Decreasing the signal length did not have an effect on dMI-based PAC estimates. When using the MI to estimate PAC, small fluctuations were observed as an effect of signal length. MVL and PLV appeared to be most affected by signal length, especially for the low frequency signals of 2 Hz and 5 Hz. The erroneously elevated PAC values of MI and PLV were also present in the analysis across different signal lengths.

## Discussion

The current work presents the dMI as a novel measure of PAC. Our dMI has been designed to be easily interpretable on a stationary interval between 0 and +1, and specific to the frequency of interest only. The performance of dMI, PLV, MVL, and MI was investigated using artificial data under increasing levels of noise and with decreasing amounts of data. The results indicate that dMI is more robust towards varying levels of Gaussian noise and short signal durations than the other PAC methods investigated in the scope of our evaluations.

First, one characteristic of is specific to dMI in comparison to the other investigated methods is that its PAC estimates are bound to a stationary interval between 0 and 1. Bounding the output to a specific, stationary interval allows for the interpretation of absolute changes in PAC. This, in turn, facilitates the calculation of meaningful effect sizes in statistical investigations. A combination of these two pieces of information is essential for any meaningful interpretation and for the discussion of any results (Lakens, 2013; Stankovski et al., 2017).

Second, dMI was found to be highly resilient towards high levels of Gaussian noise and performed well with decreasing amounts of data within the extent of the current investigations. By contrast, PAC estimates from MI, PLV, and MVL strongly deteriorated as the levels of noise increased. While the decreasing signal duration had no effect on dMI, for PLV it led to a high number of erroneously elevated PAC estimates at lower frequencies.

Finally, in our investigations, we observed that MI tends to systematically overestimate PAC values for frequencies below the target frequency. This may be related to the scoring mechanism of MI, which calculates the entropy of the phase-amplitude histogram. This entropy was still high when low levels of Gaussian noise were added. MVL-based PAC estimates also showed poor performance at frequencies lower than the target frequency, in particular for shorter signal lengths, as it returned a high number of erroneously elevated PAC estimates within that range. On the other hand, both PLV and MVL tended to systematically overestimate PAC values at the harmonics of the target frequency. These observations underline the sensitivity of PAC estimates, and the need for a reliable solution. However, it is important to bear in mind that these observations are potentially biased, as the results are derived from artificially created data.

The current implementation of dMI assumes a sinusoidal distribution of amplitudes across the individual phase bins. This assumption is likely to hold, provided a sufficiently large sample size with independent measurements is available (Nixon et al., 2010). Our implementation of dMI also enables the user to easily visualize the phase-amplitude histogram to understand the shape of the PAC fit. In the event that the observed histogram is not Gaussian, the user may conveniently define another function for the shape for the line-fitting.

## Conclusions

Here, we presented dMI as a new measure to estimate PAC of neurophysiological data on the basis of a sinusoidal fit of the phase-amplitude histogram between two signals. The dMI has been designed to be resistant against varying levels of noise, to perform well with short signal durations, and to be easily interpretable due to the absolute boundary values of 0 and +1. Furthermore, through configurations of the parameters and/or changing of the sinusoidal scoring function, dMI is easily adaptable to the question at hand. We used simulated data to show that dMI provides a more reliable estimate of PAC than a number of other established measures. This novel measure may therefore provide a useful tool for the investigation of brain dynamics with implications for basic and clinical science.

## Abbreviations

dMI: direct modulation index
EEG: electroencephalography
MI: modulation index
MVL: mean vector length
PAC: phase-amplitude coupling
PLV: phase-locking value

## Acknowledgements

This work was supported by the German Federal Ministry of Education and Research (BMBF). We acknowledge support by the Open Access Publishing Fund of the University of Tübingen.

## Code and data availability

The code of this analysis is available at https://github.com/VoodooCode14/dmi. The implementation of dMI is available at https://github.com/neurophysiological-analysis/FiNN.

## Credit authorship contribution statement

**Maximilian Scherer**: Conceptualization, Methodology, Software, Writing – original draft, Writing – review & editing.

**Tianlu Wang**: Writing – original draft, Writing – review & editing.

**Robert Guggenberger**: Conceptualization, Methodology, Writing – review & editing.

**Luka Milosevic**: Conceptualization, Methodology, Writing – review & editing.

**Alireza Gharabaghi**: Conceptualization, Writing – review & editing, Funding acquisition.

## Declarations of competing interests

The authors declare no conflict of interest.

## Notes

https://github.com/VoodooCode14/dmi

### Competing Interest Statement

The authors have declared no competing interest.

